# Mastering the Manu – How humans create large splashes

**DOI:** 10.1101/2024.12.03.626658

**Authors:** Pankaj Rohilla, Daehyun Choi, Halley Wallace, Kai Lauren Yung, Juhi Deora, Atharva Lele, Saad Bhamla

## Abstract

Manu jumping, a popular water diving style amongst Māori people in New Zealand, focuses on creating large splashes. Divers perform aerial maneuvers such as the “utkatasana” pose, entering the water in a V-shape, and executing underwater maneuvers to maximize the splash size. Our study explores the underlying fluid dynamics of Manu jumping and demonstrates how two key parameters, the V-angle and the timing of body opening, can maximize the Worthington jet formation. To accurately replicate human manu jumping, we studied water entry of both passive solid objects with varying V angles and an active body opening robot (Manubot). The analysis revealed that a 45-degree V angle is optimal for maximizing Worthington jet formation, consistent with human diving data. This angle balances a large cavity size and a deep pinch-off depth. The body opening within a timing window of 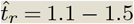 synchronizes the robot’s potential energies to be timely transferred to the cavity formation, producing the strongest and most vertical, i.e., ideal, Worthington jets. Based on our experimental findings, we propose optimal parameters for generating the largest Manu splashes. These insights offer engineering perspectives on how to modulate underwater cavity dynamics using both passive and active body formations.

## 1. Introduction

Māori Kiwis engage in their beloved traditional water sport called Manu jumping, where participants leap from bridges, docks, waterholes, and diving platforms, aiming to produce the largest possible splashes. For the Māori people, Manu jumping is more than just a recreational activity, it is a cultural way of life. Despite their mastery of this art form, the fluid dynamics underlying this unique form of jumping has not been studied before. Competitions such as the Z Manu World Champs, held across New Zealand, evaluate Manu jumping performances primarily based on the size of the splash created [1]. In Manu jumping, the act of generating a water splash is colloquially known as “popping a Manu” [2].

The hydrodynamics of Manu jumping is closely related to the water entry of the projectiles. When a projectile impacts a liquid surface, it creates an air cavity in its wake that collapses to produce a liquid jet, known as the Worthington jet [3–7]. The cavity pinch-off forms a base region that feeds and dictates the jet strength [4, 8]. The formation of Worthington jet is driven by the kinetic energy distributed across the collapsing cavity wall, with its acceleration primarily provided by the large vertical momentum around the jet base rather than the pinch-off singularity [8]. While the water entry of solid projectiles and droplets has been extensively studied [9, 10], most research has focused on minimizing splash formation and reducing damage to solid projectiles, ships and seaplanes [3, 11–14]. A study focusing on creating large Worthington splashes is still missing.

Prior studies investigating humans and animals diving into water primarily focused on understanding the safety limits and minimizing splash created by their water entry, as seen in activities like Olympic diving and birds hunting in open waters [15–22]. Additionally, post-entry active movements by divers have been investigated via physical models to minimize splash formation, commonly referred to as a ‘rip’ entry [21]. While takeoff heights in professional water sports can reach up to 27 meters, with an injury rate of 9.7 per hour of exposure, Manu jumping takeoff heights reaches up to 10 meters, which can still pose injury risks to both elite athletes and young divers [23–25]. Unlike professional diving, where the focus is on clean, head-first entries, Manu jumping involves participants landing on their backs and glutes, forming a distinctive V-shape at water entry with their bodies.

The key dimensionless numbers governing water entry dynamics of humans and solid projectiles include the Froude number, 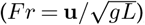, Bond number (*Bo* = *ρgL*^2^*/σ*), Weber number (*We* = *ρv*^2^*L/σ*), and Reynolds number, (*Re* = *Luρ/μ*); where **u** is the projectile impact speed on water, *L* is the characteristic length of the projectile (*L* = *W*, the width of human jumpers, *ρ* is water density, *μ* is the water viscosity, *σ* is water surface tension, and *g* is the acceleration due to gravity [5, 18, 26]. For human water entry, these numbers typically fall within the ranges 10^4^ *< Bo <* 10^5^, 1 *< Fr <* 100, 10^6^ *< Re <* 10^7^, and 10^4^ *< We <* 10^7^. These values indicate that the inertial forces of humans entering the water dominate gravitational, surface tension, and viscous forces, creating an air cavity that collapses via gravitational forces to generate Worthington jets and splashes.

Here, we investigate the fluid dynamics of Manu jumping through human data, passive and active robotic projectiles. We drop these solid projectiles in water to study the role of jumping height, V-angle during water entry, and the effect of underwater body expansion on the resulting Worthington splash. To evaluate the role of human dynamics in popping a big Manu splash, we designed and tested the water entry of solid projectiles with varying V-angles and underwater maneuvers of an active robot simulating Manu jumpers. Our work offer scientific insights in recreational water jumping sports and underscores the importance of V-shape and underwater maneuvers to create large splashes in water.

## 2. Methods

### (a) Human data collection

Human Manu jumping data was extracted from 50 YouTube videos. The body dimensions of the jumpers, including height (*H*) and width (*W*), were measured using ImageJ. The key parameters such as the jumping height (*H*_*jump*_), splash height (*H*_*splash*_), and splash speed (*v*_*j*_), were also measured using ImageJ. To avoid the scale approximation of human bodies (assumed *h* = 1.71 m), we normalized the jump height and splash height by the body width, resulting in dimensionless jump height (*β*_*jump*_ = *H*_*jump*_*/W*) and dimensionless splash height (*β*_*splash*_ = *H*_*splash*_*/W*).

### (b) Solid bodies

Five solid bodies of varying shapes, each with distinct V-angles (0^°^, 45^°^, 90^°^, 120^°^ and 180^°^), were designed to mimic the body posture of human Manu jumpers during water entry. These solid bodies were fabricated using a 3D printer (Bambu Lab X1-Carbon) with Polylactic acid (PLA) filament with 100% infill density, ensuring high-density prints for complete submersion in water. The body density and aspect ratio (*W* : *H*) of the solid bodies were kept constant at 1.24 kg/m^3^ and 1 : 4, respectively. The slamming curvatures at the apex of the solid bodies were kept consistent. A flat-head hex bolt was embedded at the center of the solid bodies, allowing them to be secured to an electromagnet. The bodies were suspended by the electromagnet at an initial height *H*_*o*_ above water, contained in a water tank of the dimensions 600 *×* 400 *×* 400 *mm*^3^ (figure A.1 in the Appendix). The Manubot was held using two electromagnets. The solid bodies were released from three different heights (*H*_*o*_), resulting in three impact speeds, estimated as 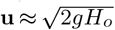, ranging from 0.75-2 m/s.

### (c) Manubot

The Manubot is designed to simulate the underwater opening dynamics of human bodies during the Manu jump, focusing on the initial water entry angle, actuation, and timing of opening. Its 3D-printed body, fabricated using the same method as described for solid bodies earlier, includes a hinge mechanism, enabling it to open actively (mimicking the underwater roll back and kick motion of human); this mechanism includes a geared motor and the angle restrictor attachments. The angle restrictors prevent the body from closing completely, maintaining a specific angle that can be adjusted by changing the length of the attachment. Figure 6.a shows the robot in its closed (top) and open (bottom) positions. The total mass of the Manubot was 115 g.

The opening of the robot body is driven by a spring-and-release mechanism, with metal nuts attached to it ends to facilitate its connection with the electromagnet system, ensuring its controlled release during water drop experiments. Inside the body, a geared motor (CL 1.5 Micro Motor) operates the release mechanism, using thread to control movement. A microprocessor (Arduino Nano) controls the timing of the release mechanism (*t*_*r*_ =0.04 - 0.32s where *t*_*r*_ denotes the body opening time), which is powered by the motor. For the no-opening case (when the Manubot never opens), the body opening time is expressed as *t*_*r*_ → ∞.

A simple spring-mass model was used to measure the spring coefficient (*k*) of each elastic band used in the Manubot. The experimental setup involved suspending the elastic bands from a fixed support at one end and then applying known weights to the other end to measure the spring coefficient. An analytical balance (ME204TE, Mettler Toledo) was used to determine the object’s mass. The change in elongation (*ΔL*) of the elastic bands was measured using digital calipers. Three ring-shaped elastic bands were tested, with diameters of 9.5 mm, 7.9 mm, 25.4 mm, respectively. Using Hooke’s Law (*k* = *F/ΔL*), the spring coefficients were calculated as k = 59.6, 80.8, and 134.2 N/m, corresponding to the most flexible, intermediate flexible, and less flexible rubber bands, respectively.

### (d) Imaging and image processing

A high-speed camera (Photron FASTCAM MINI AX, frame rate = 2.000 fps) mounted with a Nikon Zoom lens (70-200mm f/2.8G ED VR II AF-S, Nikkor) was used to capture the impact of solid bodies on water, along with the resulting cavity formation and Worthington jet dynamics. The water tank was illuminated from two angles, with a light diffuser attached to the water tank.

The kinematics and angular opening dynamics of the projectile were measured by tracking its apex while for the Manubot, three points were tracked, two at the edges and one at the apex (or center) of the solid bodies. We used the Photron Fastcam Analysis (PFA), Photron Fastcam Viewer (PFV4), and DLTdv8 (a MATLAB-based digitizing tool) to track the coordinates of these points frame-by-frame. To quantify the Worthington jet, we measured the splash height in PFV4 and the jet area using a custom MATLAB script. The splash area was defined as the Worthington jet region above *y >* 0.1*L*_*m*_, where *L*_*m*_ is the Manubot’s single arm length. Images were binarized with specific threshold to capture the edge of Worthington jet. The inner area of jet was filled with region identification algorithm and measured across all time steps.

## 3. Results and Discussion

### (a) Manu jumping in humans

#### (i) Aerial maneuvers and the V-shape entry

A Manu jump comprise of four distinct stages: water entry in a V-formation, a rollback and kick motion underwater to enlarge the air cavity, the closure and collapse of the air cavity, and the formation of a Worthington splash. Figure 1.a shows an overlay of sequential snapshots illustrating the step-by-step aerial maneuvers of an individual performing a Manu jump from a diving board into a swimming pool. To gain momentum, the individual pushes the diving board downward (steps 1–2) before jumping vertically into the water executing an ‘utkatasana’ pose (steps 3–4). During vertical free fall, the individual folds their body into an L-shape (step 5) before water entry while transitioning to a V-shape upon water entry (step 6).

**Figure 1.**
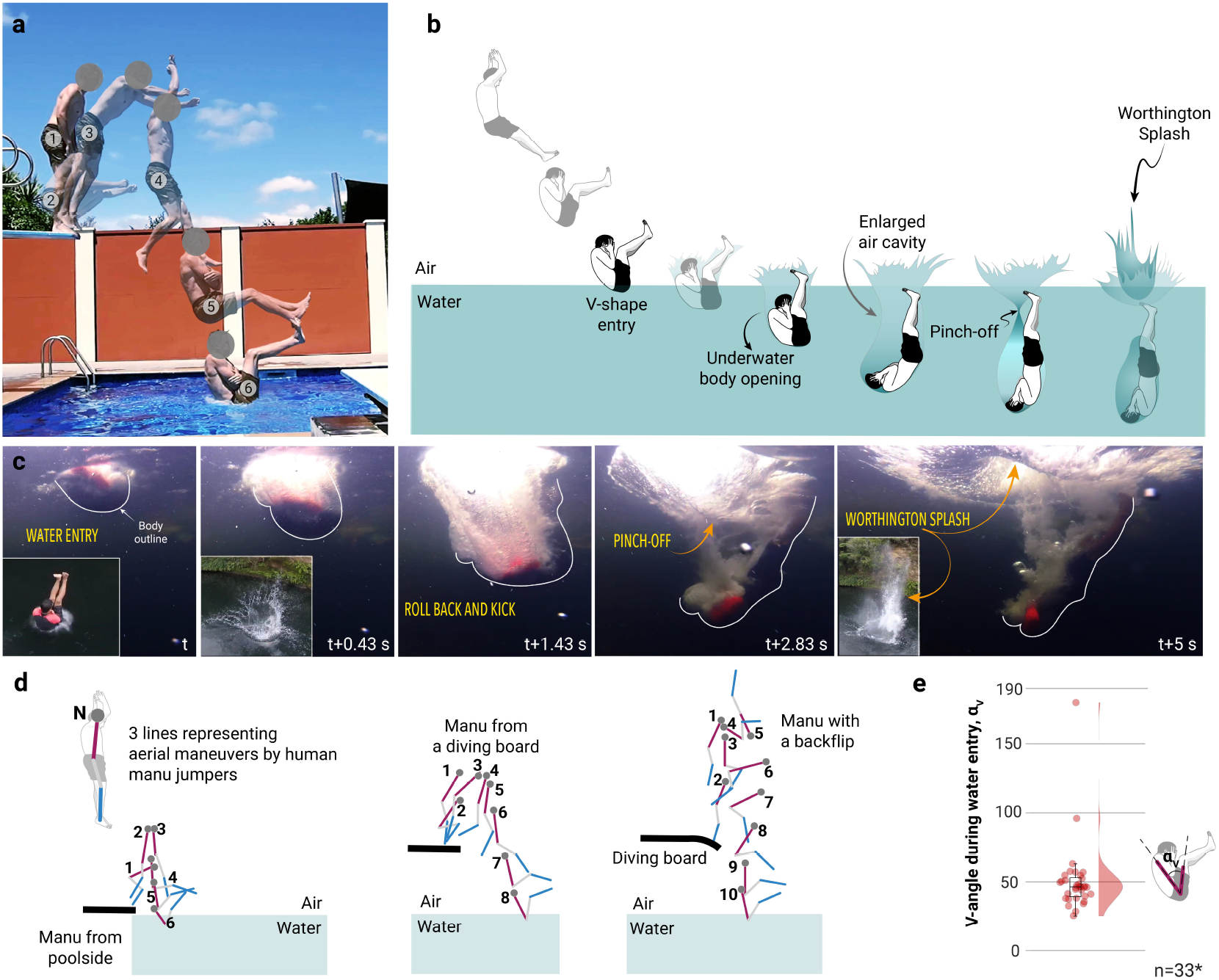
**(a)**. A composite image showing a human performing a Manu jump, illustrating the successive stages of aerial maneuvers, leading to a V-shaped entry into the water (Video credits: Bradford | Youtube). **b)** Illustration describing a human performing a Manu jump. **c)** Water entry and subsequent opening of human underwater to enhance the size of the air cavity to create a Worthington splash (Video credits: Bomb School | Youtube). **d)** Different types of aerial maneuvers to form a V-shape at water entry. **e)** V-angle formed by humans entering water to pop a Manu splash. Median V-angle, *α*_*v*_ = 46.36 with a standard deviation of ∼ 26.29 (n = 33).

A more holistic depiction of a Manu jump covering both aerial and underwater maneuvers to create a large Worthington splash is presented in Figure 1.b. In addition to executing the aerial transitions into a V-shape at water entry, Manu jumpers perform underwater maneuvers. These include a rollback and kick motion, where they extend their bodies underwater - moving their head and back downwards while feet point upwards. This movement expands the air cavity, which eventually reaches its maximum size before collapsing and closing at a pinch-off point, forcing water upwards in form of a Worthington splash. Figure 1.c shows a sequence of snapshots of an individual performing an underwater body expansion. Before entering the water, Manu jumpers bend their body while keeping their back straight, forming a V-shape. This water entry creates a crown splash (figure 2.d, inset of the second snapshot), deforming the water surface and initiating the formation of an air cavity. If the body remains compact, the cavity will likely close and collapse. However, Manu jumpers counteract this by extending their body underwater, rolling backward and kicking downward to enlarge the air cavity (third snapshot). Eventually, the cavity collapses and pinches off (fourth snapshot), forcing the liquid rapidly upward to create a splash, as shown in the final snapshot inset of figure 2.d.

**Figure 2.**
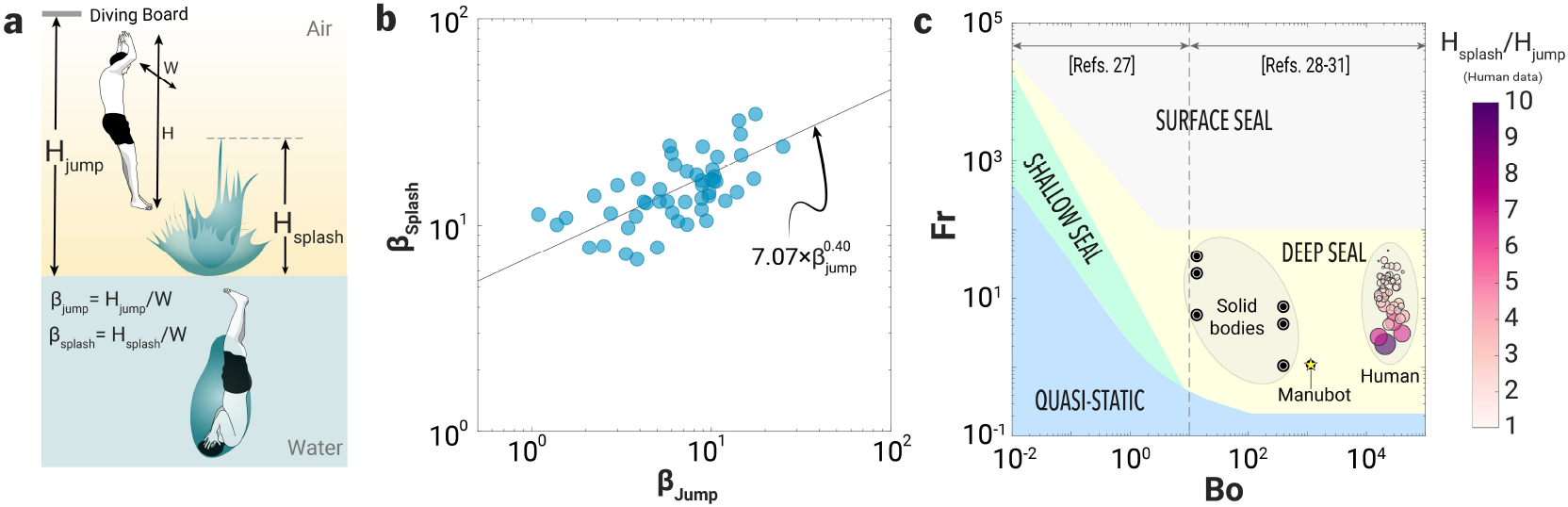
Effect of jumping height and underwater dynamics on the splash height in Manu jumping. **a)** Jumping and splash parameters illustrated for a Manu jump. **b)** Dimensionless splash height (*β*_*splash*_) vs. dimensionless jumping height (*β*_*jump*_), n = 50. Solid line represents a power law fit to the Manu jumping data (coefficient of determination, *R*^2^ ≈ 0.41). **c)** Regime map for the cavity shape created by the water entry of solid spheres, solid shapes [27–31], humans performing Manu jumps, and the Manubot.

From human jumping video observations, we hypothesize that the underwater maneuvers performed by Manu jumpers not only enlarge the air cavity but also delay its pinch-off, enabling stronger gravitational forces to retract the deformed water surface and generate a larger Worthington splash. These underwater movements are thus crucial in popping a big Manu splash, particularly when other factors like jumping height and body dimensions are similar.

Manu jumping, as a recreational sport, invites creativity and personal flair, with individuals showcasing aerial maneuvers like backflips and diving from various platforms such as docks, cliffs, trees, diving boards, and pool edges. Figure 1.d illustrates the body orientation of Manu jumpers during various types of Manu jumps, based on frames extracted from YouTube videos. In the first sequence, a jumper performs a Manu jump from a shallow height (poolside), forming the V-shape quickly due to the limited fall distance. In the second sequence, corresponding to the jump shown in figure 1.a, the individual leaps from a diving board, descends vertically, and fold their legs to form a V-shape before water entry. In the third instance, the jumper uses a diving board to generate upward momentum for a backflip, skillfully controlling their orientation to achieve a V-shape upon water entry (figure 1.d). Despite differences in height and aerial maneuvers, all jumps consistently result in a V-shape at water entry. The median V-angle during water entry of human Manu jumpers was ∼ 46.36^°^ (figure 1.e).

#### (ii) Human kinematics in Manu jumping

Apart from entering the water in a V-formation, the underwater maneuvers play a key role in generating a Manu splash. When humans enter the water, their impact deforms the surface, creating a large air cavity. In this process, the inertial forces of the body temporarily overcomes the gravitational forces that maintains the water surface (*Fr >* 1). As the gravitational pull of the water overcomes the inertia of the human body, the air cavity collapses and pinches off. This collapse rapidly displaces water upward from the pinch-off point, driving the formation of an accelerating jet or a splash that ascends until surface tension and gravity dominate, decelerating its ascent. Eventually, the jet or splash halts and falls back into the water. For human data, we studied the effect of varying the take-off height (*H*_*jump*_) on the splash height (*H*_*splash*_).

Increasing the height of the jump increases the impact speed (**u**) of the Manu jumper on water surface, i.e., 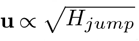. In Manu jumping championships, the maximum jumping height is limited at 10 meters to ensure the safety of participants. While jumping height is a critical factor, participants’ body size and underwater maneuvers also significantly influence the resulting splash height. However, since the YouTube data used in our analysis lack information on body weight, our study focuses on the effect of jumping height on splash dynamics. We examined the effect of dimensionless jumping height (*β*_*jump*_ = *H*_*jump*_*/W*, where *W* is the width of the Manu jumper) on the dimensionless height (*β*_*splash*_ = *H*_*splash*_*/W*) of the resulting Worthington jets (see definition of these dimensions in figure 2.a). The dimensionless splash height increased with jumping height as 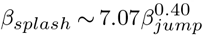 (figure 2.b). The cavity pinch-off dynamics in human Manu jumps can also be characterized using Froude number (*Fr*) and Bond number (*Bo*) plots [28]. For similar *Bo*, lower *Fr* rendered higher *H*_*splash*_*/H*_*jump*_ ratios which decreased with increasing *Fr*. This suggests that for increasing *Fr*(∝ **u**), *H*_*splash*_ does not increase linearly with increasing *H*_*jump*_. Human water entry during Manu jumps falls within the deep seal regime, specifically in the range of 10^4^ *< Bo <* 10^5^ and 1 *< Fr <* 100 (figure 2.c). This indicates that the inertial forces of the human body entering the water during a Manu jump, dominates the surface tension and gravitational forces. Similarly, the other shapes used in this study also fall within the deep seal regime, highlighting consistent dynamics across different configurations of passive and active projectiles.

### (b) Splash dynamics of solid projectiles

#### (i) Water entry, cavity formation and the Worthington splash

We visualized the water entry of V-shaped projectiles (or wedges) at varying impact speeds to qualitatively compare their entry dynamics, focusing on the formation of air cavities that close and collapse to produce Worthington splashes. We studied the water entry dynamics of projectile shapes with V-angles of 45^°^, 90^°^, and 120^°^ (figure 3.a) - investigating cavity formation and Worthington splash characteristics to elucidate the role of V-angle in human Manu jumping. Additionally, we used a flat vertical shape (*α*_*v*_ = 0^°^) simulating a human folding their body to align their back and legs parallelly upwards, and a flat horizontal projectile representing the scenario of a human landing on water flat on their backside (*α*_*v*_ = 180^°^).

**Figure 3.**
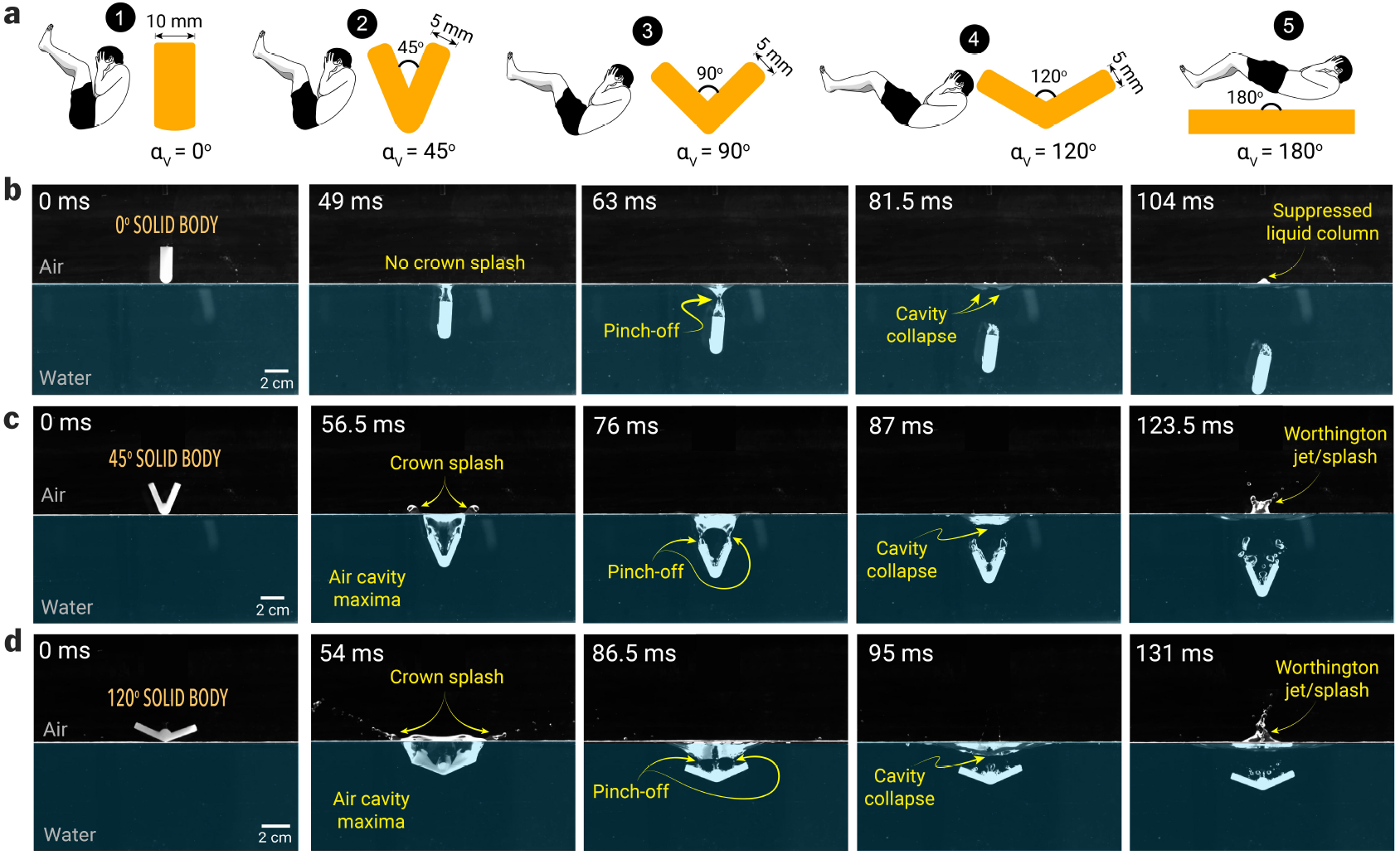
Water entry dynamics of solid bodies with varying V-angles (*α*_*V*_). **a)** Solid bodies and their corresponding human formation during water entry in our study. Human bodies are ≈ 𝒪(10^2^) times larger than solid bodies. Sequence snapshots of: **b)** a straight solid body (*α*_*V*_ = 0^°^) entering water at an impact speed of 0.75 m/s which generates a small air cavity and a shallow pinch-off point, forming a suppressed liquid column, **c)** a solid body with *α*_*V*_ = 45^°^ entering the water at a speed of 2 m/s, generates a larger air cavity that pinches off at two points, resulting in a bifurcated Worthington splash, and **d)** a solid body with *α*_*V*_ = 120^°^, entering the water at a speed of 1.5 m/s, generating a wider air cavity forming a single pinch-off point, resulting in a focused Worthington jet.

Here, we present sequential snapshots of water entry and Worthington jet formation for three representative cases with *α*_*v*_ = 0^°^, 45^°^, and 120^°^, representing the most commonly observed conditions in human water entry. These projectiles generated a range of Worthington splashes, from the smallest to the largest, resulting from cavity collapse after their water entry at varying impact speeds.

*Case i):* A solid projectile with *α*_*v*_ = 0^°^ impacts the water at a speed of 0.75 m/s. Due to its elongated shape, the width of the air cavity formed closely matches the width of the projectile itself and collapses almost immediately after the body is fully submerged (*t = 49 ms*), resulting in minimal to no crown splash. After the air cavity pinches off, it collapses (*t = 81*.*5 ms*), displacing water upward to form a suppressed liquid column, with its maximum height captured in the final snapshot (*t = 104 ms*). In competitive diving, athletes often adopt this shape upon entry and quickly transition to an L-shape underwater to ensure the cavity collapses as close to the surface as possible, minimizing or eliminating splashes.

*Case ii):* In another representative sequence of snapshots showing the water entry of solid projectiles, a body with a *α*_*v*_ = 45^°^ impacts the water at a speed of 2 m/s, creating a crown splash (*t = 56*.*5 ms*) and trapping a large volume of air compared to the first case. Notably, as the body descends further into the water, the air cavity closes, forming two distinct pinch-off points (*t = 76 ms*) at the trailing edges of the projectile. As the air cavity pinches off, two jets emerge from the pinch off location in upwards direction (*t = 87 ms*) which merge to form a forked Worthington splash (*t = 123*.*5 ms*). This case represents the majority of the water-entry cases of the Manu jumpers where the median angle of water entry is ∼ 48^°^. It is important to note that in human Manu jumping, the human body and the resulting air cavity formed in the water are not symmetrical, unlike the projectiles studied. This asymmetry leads to the formation of irregular, asymmetric Worthington splashes (last snapshot in figure 1.c).

*Case iii):* The third representative case of Manu jumping involves the water entry of the solid projectile with a 120^°^ V-angles solid projectile impacting water at **u** = 1.5 m/s (figure 3.d). The water entry of this shape created a larger air cavity (*t = 54 ms*) in water accompanied by a wide crown splash in comparison to the prior cases. The proximity of the trailing edges of the V-shaped projectile and the bolt attached at its apex to secure it to the electromagnet interferes with the air cavity’s pinch-off dynamics. This interference results in three pinch-off points (*t = 86*.*5 ms*) merging into a single upward-moving fluid curvature (*t = 95 ms*). This cavity eventually collapses, producing a single jet-like splash (*t = 131 ms*). Additionally, the 120^°^ configuration generates two flat water curtains on either side of the shape, merging to form a single splash (SI video 1).

The impact speed (**u**) and the V-angle of the solid projectile shapes affects the air cavity pinch-off depth (*H*_*po*_) and the projected area of the cavity at the pinch-off (*A*_*c*,*po*_); these parameters collectively affect the Worthington splash outcomes, quantified in terms of the maximum splash height (*H*_*max*_) and the maximum Worthington splash area (*A*_*s*,*max*_). Various stages of a solid projectile entering water, creating an air cavity and a crown splash, and finally a Worthington splash following the air cavity pinch-off are illustrated in figure 4.a. The average pinch-off depth of air cavities increased with decreasing *α*_*v*_ from 120^°^ to 45^°^ (figure 4.b), whereas the pinch-off depth was similar for 180^°^ and 120^°^ shapes. For the 0^°^ solid projectile entering water at **u** = 1.5 − 2 m/s, air cavity dynamics differ due to a thinner cavity formed by a single edge in its wake, leading to lower pinch-off depths compared to the 45^°^ projectile, which generates wider cavities with two edges and results in larger pinch-off depths at similar impact speeds.

**Figure 4.**
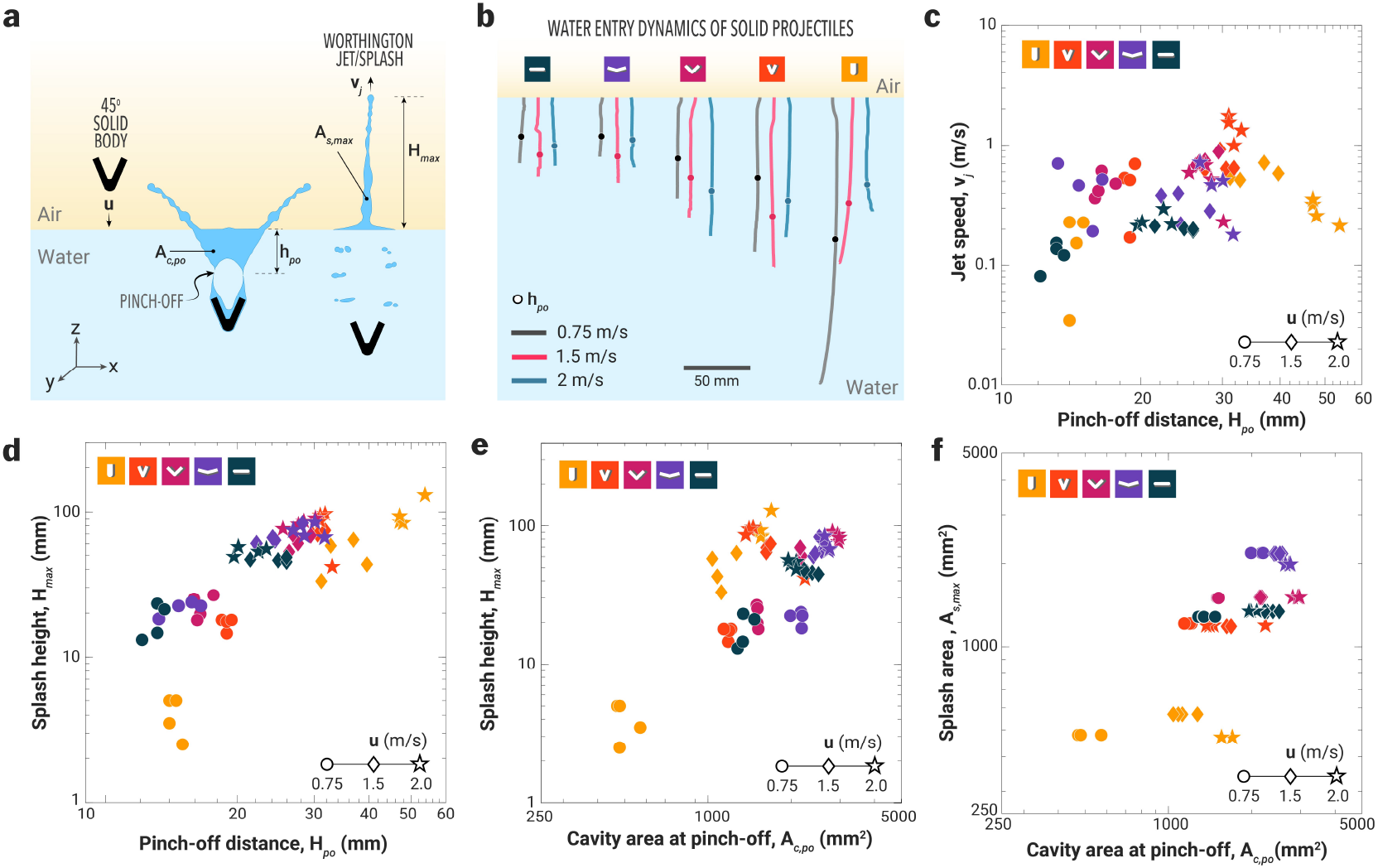
Splash dynamics of solid projectiles entering water at different impact speeds (0.75–2 m/s). **a)** Schematic showing a 45-degree-shaped projectile entering water at an impact speed of **u** m/s. The impact generates a crown splash as an air cavity forms and collapses, pinching off to produce a Worthington jet with maximum splash area *A*_*s*,*max*_ and height *H*_*max*_. **b)** trajectories and pinch-off depths of air cavities formed by various solid projectiles entering water at different impact speeds. **c)** relationship between pinch-off depth and Worthington jet speed, **d)** splash height as a function of pinch-off depth, and **e)** effect of varying pinch-off depth to enlarge the cavity area at the pinch-off point, and **f)** Worthington splash area as a function of cavity area at the pinch-off point. Note: Each data point represents a single experiment, with four replicates conducted for each experimental condition.

#### (ii) Projectiles with *α*_*v*_ = 45^°^ generate fastest Worthington jets

The 45^°^ solid projectile entering water at **u** = 2 m/s (*H*_*po*_ ∼ 30 mm) produced the fastest Worthington splash, with maximum speed reaching up to **v**_*j*_ ∼ 2 m/s (figure 4.c). Meanwhile, other angled shapes (90 and 120 for all impact speeds) and 45 shape (0.75 and 1.5 m/s) exhibited jet speeds between 0.2–1 m/s for pinch-off depths ranging from 12-30 mm. For the vertical shape (*α*_*v*_ = 0^°^), water entry produced suppressed liquid columns at lower impact speeds (**u** = 0.75 m/s) and focused tall jets at higher impact speeds (**u** = 2 m/s). The maximum jet speeds for this shape, regardless of whether the pinch-off point was shallow or deep, fell within a range of **v**_*j*_ ∼ 0.3 − 0.4 m/s. The flat shape (*α*_*v*_ = 180^°^) generated Worthington splashes with relatively slower jet speeds, ranging between 0.1–0.3 m/s across varying impact speeds.

#### (iii) The role of the pinch-off depth and the cavity size on the Worthington splash

We studied the effect of varying V-angles of solid projectiles on pinch-off depth (*H*_*po*_) and air cavity size (*A*_*c*,*po*_), and their subsequent effects on splash height (*H*_*max*_) and splash area (*A*_*s*,*max*_) (figure 4.d-e). The solid projectile with *α*_*v*_ = 0^°^ entering water at **u** = 0.75 m/s produced the shortest Worthington jet, or a suppressed liquid column (*H*_*max*_ ∼ 2.2 mm), with a pinch-off depth (*H*_*po*_) of 16 mm and the smallest cavity area (500–700 mm^2^). At a higher impact speed of 2 m/s, the same projectile shape produced the tallest Worthington jet (*H*_*max*_ ∼ 200 mm), forming a cavity with an area of approximately 16, 000 mm^2^ and pinching off at a depth of *H*_*po*_ ∼ 55 mm. Due to the thinner Worthington jets produced by this shape, their maximum projected area remained the smallest, approximately 500–600 mm^2^, across varying impact speeds (figure 4.f and to row of 5). When this shape was dropped parallel to the water surface (*α*_*v*_ = 180^°^) at a low impact speed of 0.75 m/s, the splash height ranged from 15 to 25 mm, with an air cavity area of around 1500 mm^2^ at a pinch-off depth (*H*_*po*_) of 12–14 mm. At a higher impact speed of 2 m/s, the splash height increased significantly to approximately 60 mm (with splash projected area of 1200-1300 mm^2^), with a pinch-off depth of 50–60 mm and an air cavity size (*A*_*c*,*po*_) of 2500-3000 mm^2^.

For solid projectiles with *α*_*v*_ ranging from 45-120^°^, the splash size (*H*_*max*_) increased with impact speed during water entry (figure 4.d-e), however, the splash area remained nearly same, indicating the jet became thinner and taller with increasing impact speed of the projectile. Amongst the angled shapes, the 45^°^ projectile produced the tallest splash (∼ 100 mm with *A*_*s*,*max*_ of ∼ 1200 mm^2^) corresponding to a pinch-off depth of ∼ 32 mm and a cavity area of ∼ 1200 mm^2^. At an impact speed of 1.5 m/s, the splash heights were comparable across different angled projectiles; however, the 45^°^ shape exhibited the deepest pinch-off depth and a smaller cavity area than wider angle shapes (figure 4.d-e). In Manu jumping, the air cavity created by the human body dictates Worthington jet dynamics. The 45^°^ shape demonstrates that a smaller cavity area but deeper air cavity pinch-off can generate splash heights comparable to those of wider angles (90^°^ and 120^°^), potentially reducing the risk of injury.

The snapshots of the maximum splash heights for various shapes entering water at varying impact speeds is shown in figure 5. At low impact speeds (**u** = 0.75 m/s), the splash size and shape increased with increasing V-angle, where 0^°^ shape forms a minimal splash, the 45^°^ shape forms a forked jet and the 120^°^ creates a small and thick jet. At medium impact speeds of **u** = 1.5 m/s, the solid body with 0 degree angle creates a bigger splash than its lower speed counterpart, while the Worthington jets from shapes like 45^°^ and 120^°^ cover a greater splash area. Notably, the 45^°^ shape achieves a higher splash height than other shapes at impact speed of 1.5 m/s. Further increase in impact speed to 2 m/s for 0^°^ shape resulted in tallest jets with focused tips, tall jets with a wider base for 45^°^ shape and a shallow jet with satellite drops for 120 degree shape at this highest impact speed.

**Figure 5.**
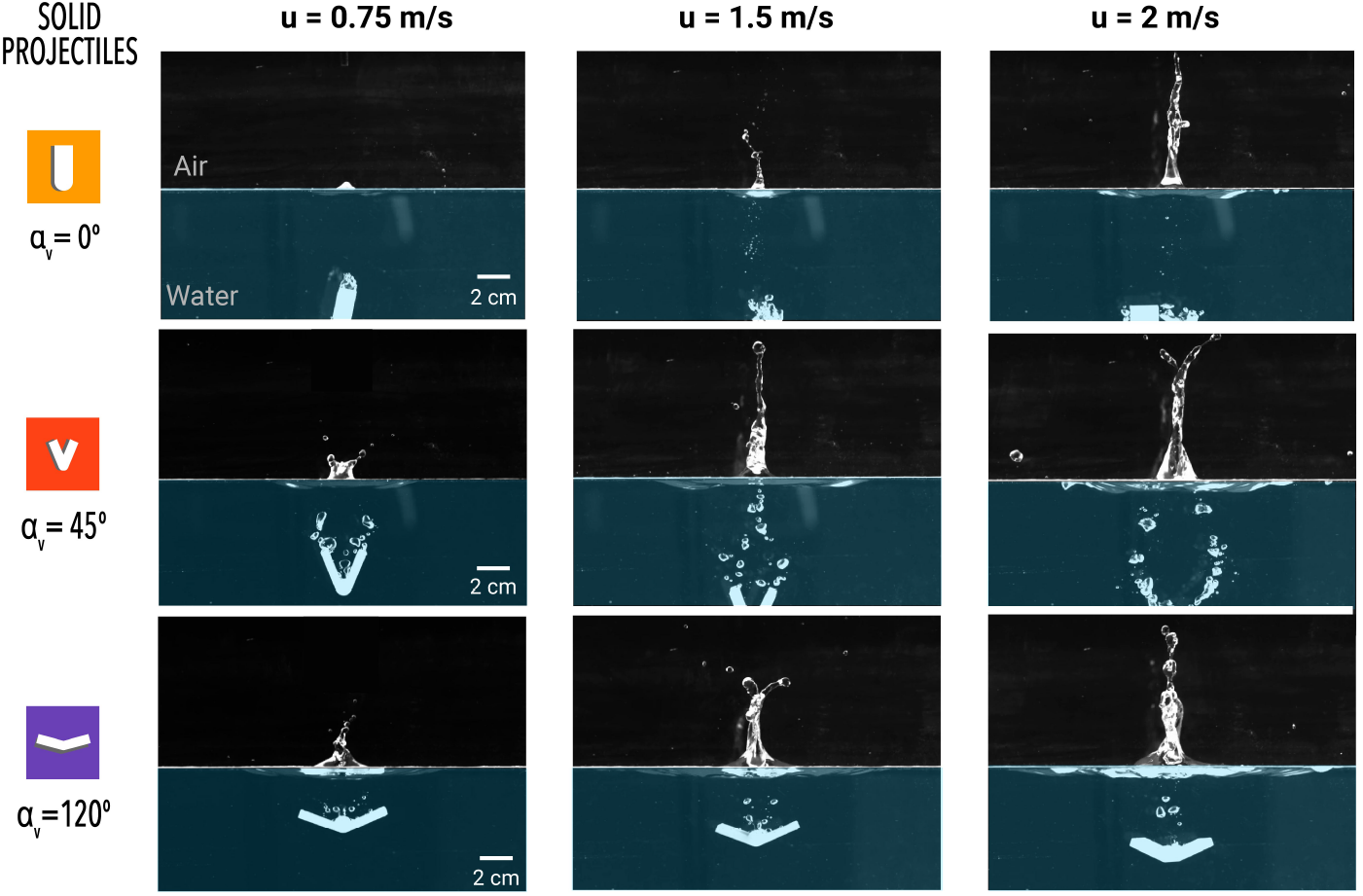
Snapshots showing the maximum height of Worthington jets achieved for water entry of solid objects with varying V-angles and impact speeds.

**Figure 6.**
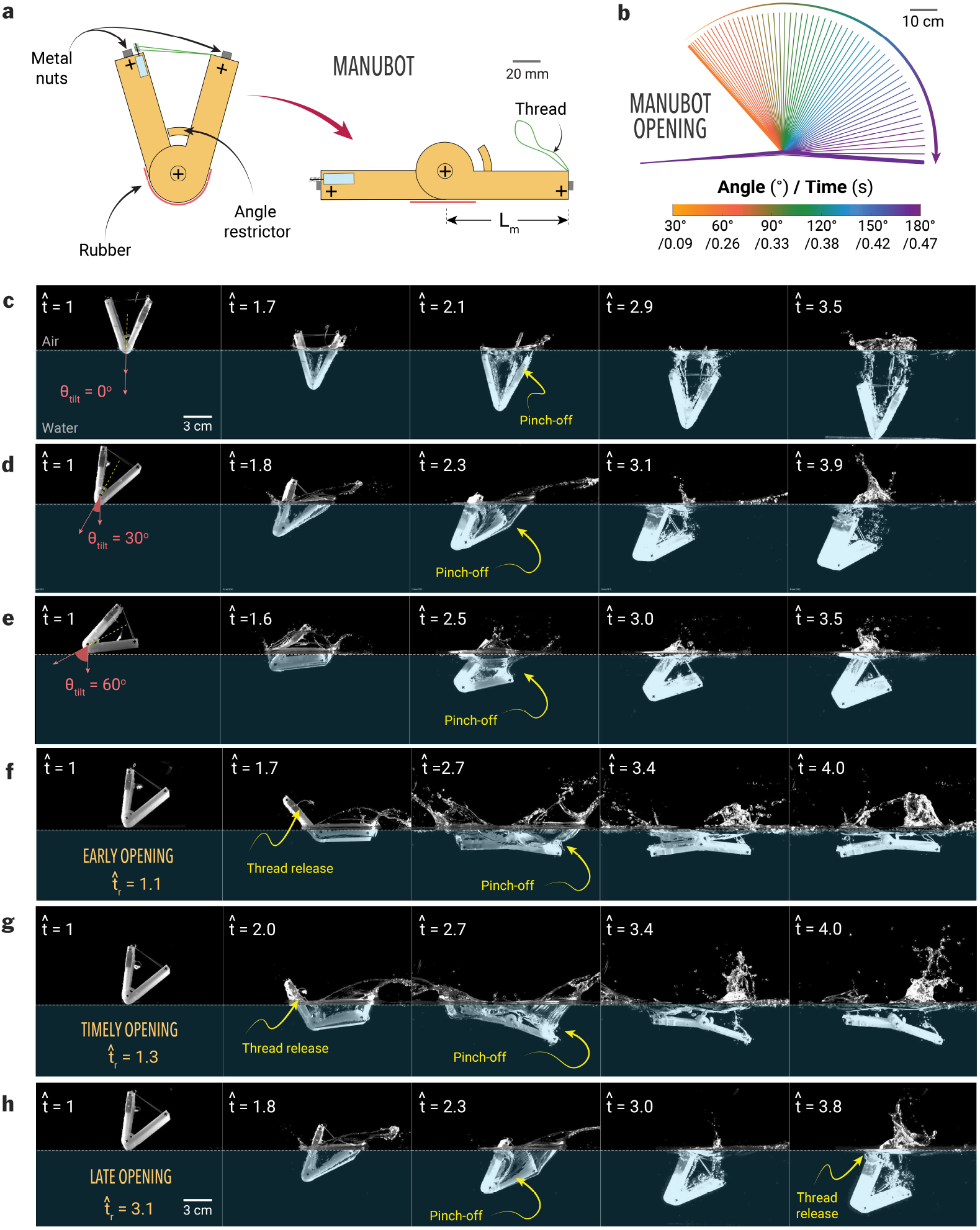
Manubot and its water entry with varying body tilt angles (*θ*_*tilt*_) and body opening times 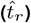. **a)** Manubot with release mechanism and expanding rubber. **b)** Speed of Manubot’s motion. **c-e)** Water entry of Manubot without body opening 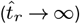 with various tilt angle of (c) *θ*_*tilt*_ = 0^°^, (d) *θ*_*tilt*_ = 30^°^, and (e) *θ*_*tilt*_ = 60^°^. **f-h)** Time sequence of Manubot falling on the water surface depending on the release time: (f) early opening 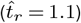, timely opening 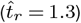, and (h) late opening 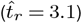. Here, the Manubot is shown from the moment it comes into contact with water at 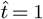, which starts to fall at 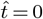.

Although V-angled projectiles offer insights on effect of V-angle and impact speed on Worthington splashes, they lack the active underwater movements of human Manu jumpers. To address this, we developed a robotic system capable of executing controlled underwater maneuvers, enabling the creation of large, asymmetric Manu splashes that closely resemble the Manu splashes generated by humans.

### (c) Manubot

#### (i) Timely body opening increases Worthington jet strength

Using high-speed imaging of the underwater body opening of the Manubot (figure 6.a) and high-speed imaging (figure A1 in the Appendix), we elucidate the the role of the onset of body opening on the Worthington jet dynamics. Here, the time (*t*) is normalized with the impact velocity (**u**) of Manubot and its arm length (*L*_*m*_), defining dimensionless time as 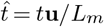. The dimensionless body opening time is defined as 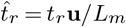, where *t*_*r*_ represents the body opening time, which can be controlled by the signal from the microprocessor. Before studying the effect of underwater opening, we investigate the role of the tilt angle of the Manubot (*θ*_tilt_, the angle between the gravity direction and the centerline of the robot) on the splash size. We determined the optimal tilt angle for water entry of the Manubot (*α*_*v*_ = 45^°^) to be 30^°^ for the splash generation (figure 6.c-e and SI video 2). At a small tilt angle (figure 6.c), the impact creates two separate air cavities 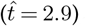, followed by a weakly developed Worthington jet 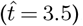. For a high tilt angle case (figure 6.d-e), a single air cavity forms on the right side of the Manubot, concentrating and strengthening the Worthington jet. However, at excessively high tilt angles (figure 6.e), the left side of the robot obstructs jet development 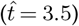. Thus, *θ*_tilt_ = 30^°^ is found to be the optimal angle for the strongest Worthington jet (figure 6.d).

After falling from the initial height (*h* ≃ *L*_*m*_ where *L*_*m*_ is the arm length of Manubot), the Manubot makes contact with water at 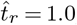, and depending on the body opening time 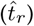, different cavity and splash dynamics occur as follows (figure 7.f-h and SI video 2):

**Figure 7.**
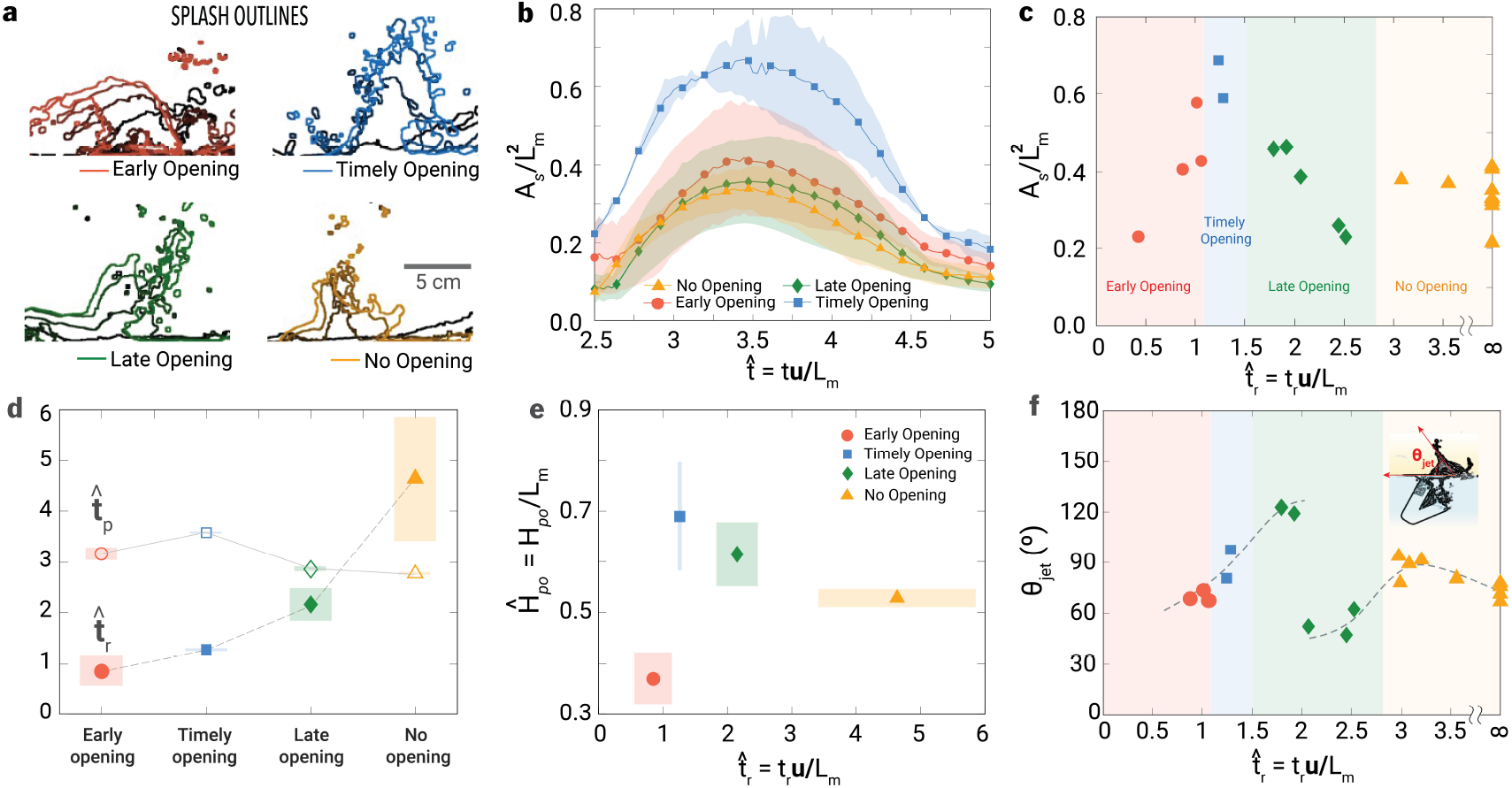
Splash and cavity dynamics of Manubot with varying body opening time 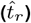. **a)** Overlay of splash development with time interval of 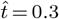, **b)** Time-varient splash area (*A*_*s*_) normalized with robot arm length 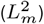, **c)** Dimensionless splash area 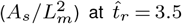, **d)** Averaged body opening time (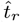, open symbols) and averaged pinch-off time (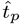, closed symbols) depending on cases, **e)** The normalized cavity depth (*Ĥ*_*po*_ = *H*_*po*_*/L*_*m*_ where *H*_*po*_ is the cavity depth) at pinch-off, and **f)** the jet direction (*θ*_jet_). Here, A total of 20 trials were conducted with the Manubot across all regimes: 4 for early opening, 2 for timely opening, 5 for late opening, and 9 for no-opening. The shade in (b), (d), and (f) indicates the error bar. 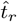 in (d) for the no-opening case exhibits a large error bar due to the absence of an upper limit.

##### Early opening 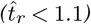

The early opening case (figure 6.f) corresponds to when the Manubot unfolds its body before water contact or at a shallow depth (*<* 0.5*L*_*m*_). The right side of the robot pushes water downward, creating large drag, while the left side remains mostly exposed to air 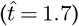. This configuration creates counterclockwise torque as two robot parts expand, and the robot eventually orients itself parallel to the water surface. Here, the accelerated left part of the robot arm creates a large splash at 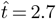. The increased contact area increases drag force, significantly decelerating the robot’s submergence. During the impact, two air cavities are formed on both the right and left sides (with the right cavity volume larger than the left) and pinch off at a shallow depth of *Ĥ*_*po*_ = *H*_*po*_*/L*_*m*_ = 0.34. The weak Worthington jets are generated on both sides due to the dissipation of the Manubot’s potential energy into two Worthington jets.

##### Timely opening 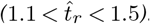

At this time range, the body opens when the robot is submerged at ∼ 0.5*L*_*m*_ at 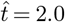. Since the left side is sufficiently submerged, the right side of robot can resist the counterclockwise torque and further pushes fluid downward, deepening the pinch-off depth to ∼ 0.7*L*_*m*_. Therefore, the formation of the air cavity and the Worthington jet is concentrated on the right side, creating the large water column on the right side of the robot.

##### Late opening 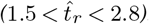

In the late opening case (figure 6.h), the body expansion starts nearly at the pinch-off time 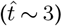. Since the air cavity has already been developed, the opening movement does not significantly affect cavity collapse or the strength of the Worthington jet. However, the jet shape is affected by the body opening time (figure 7.a), which will be discussed later.

##### No-opening 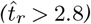

When the body opening occurs sufficiently late, its movement becomes completely irrelevant to cavity dynamics, including cavity formation and Worthington jet generation, and can be regarded as an effectively no-opening case.

#### (ii) Timely opening deepens the pinch-off and enables the vertical Worthington jet

The area of the Worthington jet has been quantified by the binarization of light intensity with certain threshold in high-speed images [32]. Figure 7.a confirms that the maximum area and height of the Worthington jet occur in the timely opening regime, while a mismatch in timing results in jet deflection in the early and late opening cases. The no-opening case produces the weakest water jet. For all times, the area of the Worthington jet (normalized by the square of one side length of the robot, 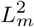) in the timely opening case outperforms the other cases which are comparable to each other (figure 7.b). Figure 7.c shows that this maximum Worthington jet area can only be achieved when the robot starts to release within a narrow range of time 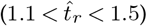, implying that accomplishing perfect Manubot performance for humans requires subtle body maneuvering.

The opening time 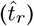 for four regimes have been identified in figure 7.d: early opening 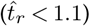, timely opening 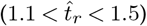, late opening 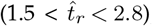, and no-opening 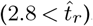. In terms of pinch-off time (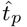, defined as 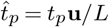 where *t*_*p*_ is pinch-off time), the air cavity pinches off at 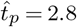 without body opening, i.e., the no-opening case, and the pinch-off is slightly delayed for all opening cases: 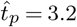 (early opening), 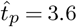 (timely opening), and 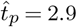 (late opening). This delay is attributed to the increased impact velocity, which creates a wider air cavity and, thereby, delays the pinch-off incident.

The timely opening case delays pinch-off the most (figure 7.d) and also achieves the greatest depth (figure 7.e), as the right side of the robot can effectively push water downward, contributing to cavity formation. For the timely opening case, the depth and time of pinch-off increase by approximately 30% and 15%, respectively, compared to the no-opening case.

The jet angle follows an interesting trend (figure 7.f): as 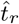 increases from 0.8 to 2, the jet angles gradually increase from 60^°^ to 120^°^, achieving vertical (90^°^) at the timely opening at 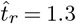. At 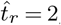, the jet direction suddenly transitions to 50^°^ and saturates at 70^°^ from 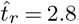. The vertical (90^°^) Worthington jet can be achieved either within a narrow range of timely openings 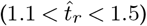 or in cases with no opening 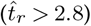.

#### (iii) Timely energy transfer enables maximizing Worthington jet

The Manubot releases each potential energies differently to its environment depending on the body opening timing (figure 8.a). The submergence depth increases as the body opening is delayed (figure 8.b) because the folded body experiences lower drag than other opening cases, allowing it to fall faster (figure 8.c). Depending on the body opening time, the slope of the body angle also differs after release (figure 8.d): timely 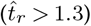 and late opening cases 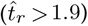 have more gradual curves than the early opening case 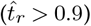. This implies that robot expansion decelerates due to drag, and momentum transfer between the robot and water is active for the timely and late opening cases.

**Figure 8.**
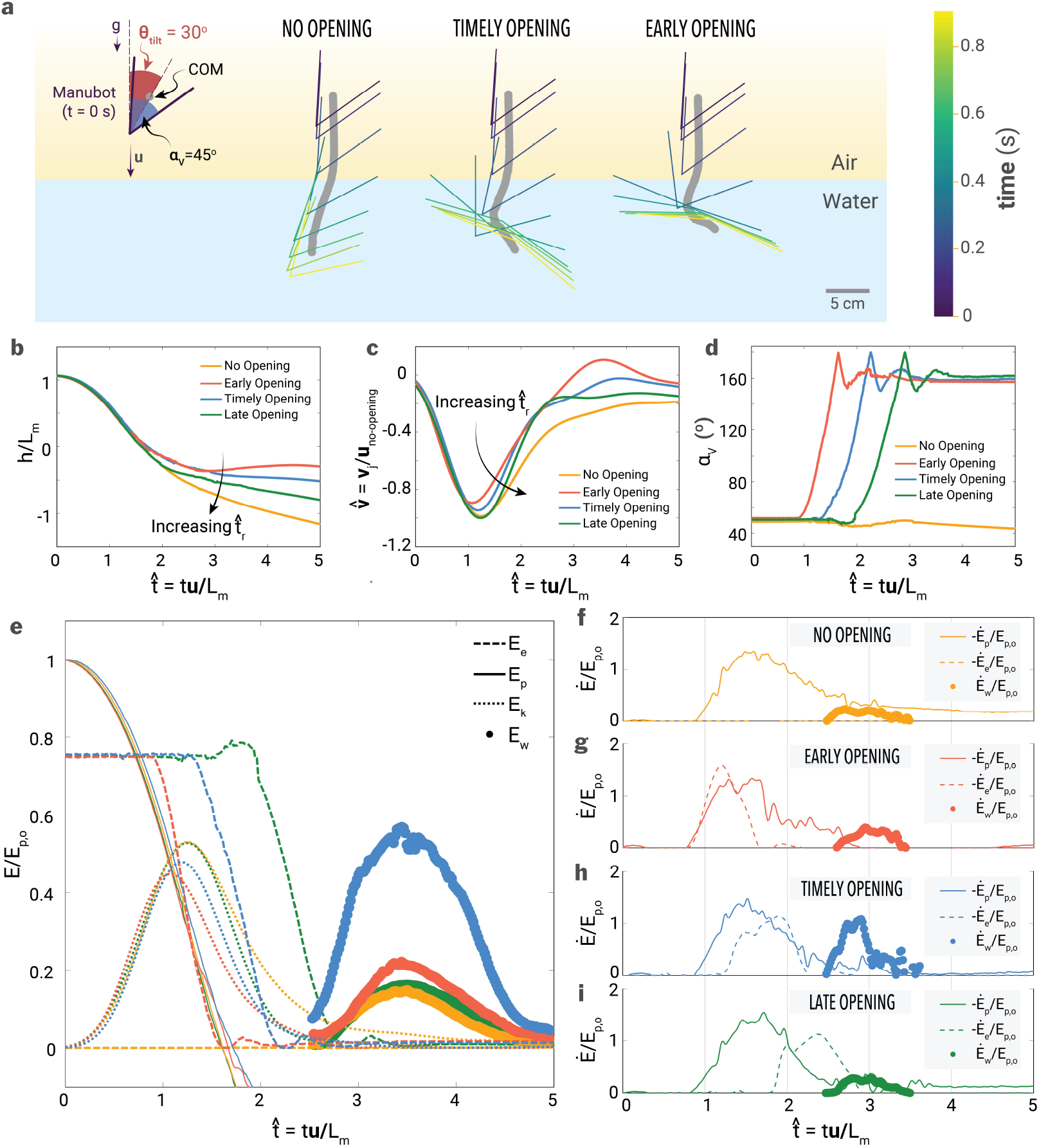
Kinematics and energy distribution of Manubot during water entry. **a)** Trajectory of the Manubot and its center of mass (COM) for scenarios with no body opening, early body opening, and late body opening inside water. The Manubot is released from air with an initial tilt angle of 30^°^. **b)** Vertical position (*h*), normalized by the Manubot length (*L*_*m*_). **c)** Vertical velocity (**u**), normalized by the impact speed in the no-opening case (*v*_*impact*_). **d)** Body angle. **e)** Energy distribution of gravitational potential energy (*E*_*p*_), elastic potential energy (*E*_*e*_), translational kinetic energy of the Manubot (*E*_*k*_), and gravitational potential energy of the water column (*E*_*w*_) for (e) early, (f) timely, and (g) late opening cases. **f-i)** Flux of mechanical (*Ė*_*p*_ + *Ė*_*k*_), elastic (*Ė*_*e*_), and jet potential energy (*Ė*_*w*_) for no-, early, timely, and late opening cases, respectively. Here, four representative cases are shown for 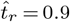 (early opening), 1.3 (timely opening), 1.8 (late opening), and ∞ (no opening). All energies and fluxes are normalized by the initial gravitational potential energy (*E*_*p*,0_).

In perspective of energy conservation, the Worthington jet is energized by two potential energies: the gravitational energy of Manubot (*E*_*p*_ = *mgh*, where *m, g*, and *h* are the mass of Manubot, gravitational acceleration, and vertical position of Manubot, respectively) and the elastic energy stored in the initially extended rubber in the robot 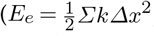, where *σk* is the sum of the spring coefficients of the rubber used in Manubot, and *Δx* is the displacement of the rubber proportional to the V-angle).

These two potential energies are transferred to the translational and rotational kinetic energy of Manubot 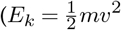, where *v* is the vertical velocity of Manubot, and 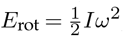, where *I* and *ω* are the moment of inertia and angular velocity of Manubot) and subsequently to the mechanical energy of water (*E*_*m*,*w*_, the sum of kinetic and gravitational potential energy of water). Some of this energy (*E*_*m*,*w*_) is temporarily stored as the gravitational potential energy of the Worthington jet (*E*_*w*_) at its highest position, and all energies are eventually dissipated as heat.

To understand how energy flows, the energy distribution has been calculated in figure 8.e. The gravitational potential energy of the Worthington jet (*E*_*w*_) is estimated from the jet area in figure 7.a by assuming it forms a circular cone with a cone angle of 45^°^ with the same cross-sectional area. Here, the rotational kinetic energy of Manubot is neglected due to the small change in tilted angle (see figure 8.a), and the air kinetic energy is neglected due to the low air density. All energy is normalized with the initial gravitational potential energy of Manubot (*E*_*p*,0_), measured at 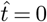. Figure 8.e shows that the gravitational potential energy (*E*_*p*_) decreases from 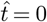 to 2 while the kinetic energy (*E*_*k*_) increases from zero and subsequently decreases after water contact 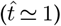, attributed to the hydrodynamic drag. For the opening cases, the elastic potential energy (*E*_*e*_) begins to decrease, releasing energy at 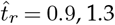, and 1.8 for early, timely, and late opening, respectively. Despite this temporal difference of *E*_*e*_, the potential energy of the Worthington jet (*E*_*w*_) for all cases emerges from 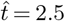 and peaks at 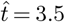, because the pinch-off time 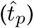 remains consistent regardless of body opening time 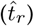, as described in figure 7.d. However, the maximum jet potential energy (*E*_*w*,max_) is significantly influenced by 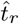: *E*_*w*,max_*/E*_*p*,0_ = 0.22 (for 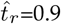, early opening case), 0.55 (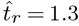, timely opening case), 0.16 (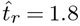, late opening case, and 0.14 (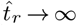, no-opening case) (figure 8.e).

Figure 8.f-i shows three energy fluxes for the mechanical energy of Manubot (*Ė*_*p*_ + *Ė*_*k*_), the elastic potential energy of Manubot (*Ė*_*e*_), and the gravitational potential energy of Worthington jet (*Ė*_*w*_). Here, all energies are normalized with the initial gravitational potential energy (*E*_*p*,0_) and the time-derivative of energy with respect to dimensionless time is expressed as 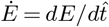. The fluxes of mechanical and elastic energy are shown in negative sign since they always decrease in this system. For all cases, the mechanical energy flux ((*Ė*_*k*_ + *Ė*_*p*_)*/E*_*p*,0_, see solid line in figure 8.f-i) emerges from the moment of water contact 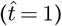 and peaks at 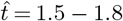 at which the center of mass passes the surface level (*h* = 0). For no-opening case (figure 8.f), the mechanical energy flux gradually decreases from its maximum and saturates to non-zero value because the Manubot keeps descending underwater due to its low hydrodynamic drag. On the other hand, for all dynamic cases in figure 8.g-i, the flux of the mechanical energy vanishes at 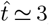, indicating that the body opening can localize the transfer of mechanical energy to water flow within short time range of 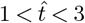 by increasing the hydrodynamic drag. Regardless of the body opening, the jet potential flux (*Ė*_*w*_ */E*_*p*,0_) begins at a similar time 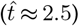, indicating that the energy transfer from the robot to the air cavity must be completed at least 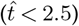. By observation, when the Manubot opens its body too early 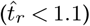, two separated air cavities are generated, weakening the Worthington jet (see figure 6.f at 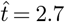 and 4.0). This suggests the lower limit of 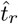 as 1.1. Since the time duration where *Ė*_*w*_ */E*_*p*,0_ *>* 0 is typically 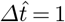 (figure 8.f-i), the optimal body opening time should be within 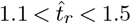, which enables two potential energies (i.e., mechanical and elastic energy) released before the Worthington jet emerges. This aligns well with the observation in figure 8.h where 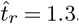. On the other hand, the Worthington jet already develops when the supply of the jet potential energy flux is being fed for the late opening cases, which proves it as off-optimal case. Therefore, the “perfect” Manu jump (i.e., optimal Worthington jet generation) requires subtle control of body opening 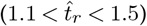 which enables successive release of mechanical and elastic energy, which can deepens the pinch-off depth and thereby results in powerful water jet.

## 4. Conclusions

Our work provides a comprehensive understanding of the fluid dynamics of Manu jumping, by integrating human data, controlled release experiments with solid V-shape projectiles, and dynamic Manubot. We show that the size of the Worthington jets or splashes, is governed by two key parameters: the V-angle and the timing of underwater body opening. These factors directly affects the dynamics of air cavity formation, pinch-off, and the resulting Worthington splash.

Human Manu jumping involves strategic aerial and underwater maneuvers to maximize air cavity size while minimizing impact forces on the body. Our analysis of 50 Manu jumping videos showed that the median V-angle during water entry was ∼ 46.36^°^, closely aligning with the optimal V-angle for solid projectiles to create large Worthington jets. To the best of our knowledge, this is the first study to investigate the cavity and splash dynamics associated with the water entry of humans and their physical models, aimed at creating large splashes. Our experiments used V-shaped projectiles, which more accurately represent the human body compared to traditional spherical or cylindrical models.

For humans, solid projectiles, and the Manubot impacting water at varying speeds, a deep-seal phenomenon occurs upon entry, where inertial forces dominate over surface tension and gravitational forces. This deep-seal regime is characterized by 1 *< Fr <* 100 and 10 *< Bo <* 10^5^, encompassing the water entry dynamics of humans, V-shaped projectiles, and the Manubot. Within this regime, increasing jumping height or impact speed of solid V-projectiles enhanced the splash size, similar to human Manu jumpers. Additionally, the Manubot effectively captured the qualitative splash dynamics of human Manu jumping by controlling the underwater body opening time.

Our experiments demonstrate V-shaped projectiles, particularly at 45^°^ angles, generate the fastest, taller and the biggest Worthington jets at higher impact speeds, emphasizing the role of cavity pinch-off depth and shape in splash generation. This supports the use of V-angle and a median angle of ∼ 46^°^ in human Manu jumps.

Our work on the dynamic Manubot highlights the role of underwater body opening in altering the the splash dynamics. Precise timing of body opening enhances the cavity depth and delays pinch-off, leading to the large Worthington splashes. Moreover, the splashes generated by the Manubot mimics the asymmetric splashes in Manu jumping, which occurs due to the asymmetric underwater body opening and non-zero tilt angles during water entry. Through the energy analysis, the synchronization between potential and kinetic energy transfer of Manubot is crucial during cavity formation for optimal splash dynamics.

Our work offers broader implications for fluid dynamics, with potential applications in naval engineering, biomechanics, and systems influenced by factors such as body elasticity, surface wettability, and environmental conditions. Additionally, this work lays the groundwork for further explorations of water entry and underwater opening dynamics in generating the Worthington splash through measurement of impact force profiles for solid projectiles, the Manubot, and human Manu jumpers. By using the Manubot and solid V-shaped projectiles, we approximated human dimensions through aspect ratio; however, future research should address projectile weight effects, an important factor in Manu jumping sports. Incorporating the anatomical complexity of human bodies into future experiments would further refine our understanding of splash dynamics. For our analysis of human Manu jumping data, we leveraged publicly available YouTube videos, which provide valuable semi-quantitative observational data. These efforts could be complemented in the future by high-precision experiments featuring divers equipped with body sensors, digital tracking markers, and high-speed imaging to simultaneously capture splash dynamics above and below the water.

Overall, our study provides a scientific understanding of the optimum V-angle of 45^°^ in Manu jumping, while emphasizing the central role of controlling underwater cavity dynamics in creating big splashes, advancing both the recreational enjoyment and competitive precision of water sports.

## A. Appendix

Figure A1 shows the experimental setup used to study the Worthington jets generated by a Manubot. The release timing of the Manubot’s two edges from the electromagnet was controlled using DC power and relays via an Arduino microprocessor, powered and programmed through a computer.

### Manubot experimental setup

**Figure A1.**
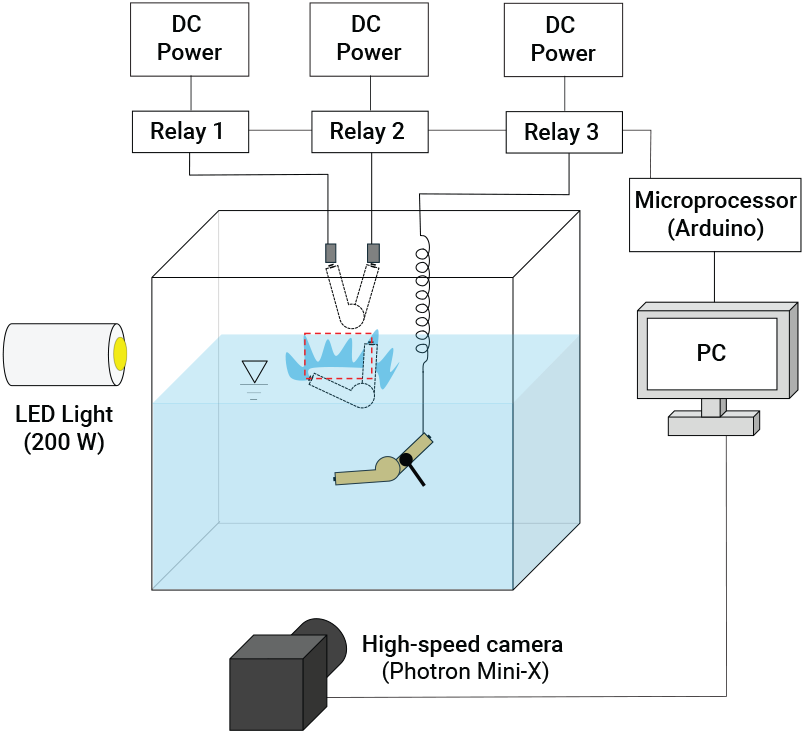
Experimental setup for studying splash dynamics of Manubot’s water entry for varying release time, opening time, and impact angles.

## Ethics

This work did not require ethical approval from a human subject or animal welfare committee.

## Data Accessibility

All data is provided in the Supplementary Information.

## Authors’ Contributions

P.R.: conceptualization, data curation, formal analysis, investigation, methodology, resources, software, validation, visualization, writing—original draft, writing—review and editing; D.C.: conceptualization, data curation, formal analysis, investigation, methodology, resources, software, validation, visualization, writing—original draft, writing—review and editing; H.W.: data curation, formal analysis, Methodology, investigation, software, visualization, writing—original draft; K.L.Y.: investigation, data curation,formal analysis, software, writing—original draft; J.D.: data curation, formal analysis, software; A.L.: formal analysis, software; S.B.: conceptualization, supervision, validation, writing—review and editing, funding acquisition, project administration; resources, investigation, methodology.

## Acknowledgements

We thank Patria Hume (University of Auckland) for insightful discussions.

## References

1 Z. Manu, Z Manu world champs, 2024.

2 Z. O’Neill, Pop some manus, diving in New Zealand with Zico O’Neill, 2023.

3 A. M. Worthington, A study of splashes (Longmans, Green, and Company, 1908).

4 S. Gekle and J. M. Gordillo, “Generation and breakup of Worthington jets after cavity collapse. Part 1. Jet formation”, Journal of fluid mechanics 663, 293–330 (2010).

5 T. T. Truscott, B. P. Epps, and J. Belden, “Water entry of projectiles”, Annual review of fluid mechanics 46, 355–378 (2014).

6 N. Kim and H. Park, “Water entry of rounded cylindrical bodies with different aspect ratios and surface conditions”, Journal of Fluid Mechanics 863, 757–788 (2019).

7 S. D. Guleria, A. Dhar, and D. V. Patil, “Experimental insights on the water entry of hydrophobic sphere”, Physics of Fluids 33 (2021).

8 S. Gekle, J. M. Gordillo, D. van der Meer, and D. Lohse, “High-speed jet formation after solid object impact”, Physical review letters 102, 034502 (2009).

9 T. Okawa, K. Kawai, K. Kubo, and S. Kitabayashi, “Fundamental characteristics of secondary drops produced by early splash during single-drop impingement onto a thick liquid film”, Experimental Thermal and Fluid Science 131, 110533 (2022).

10 T. Okawa, T. Shiraishi, and T. Mori, “Production of secondary drops during the single water drop impact onto a plane water surface”, Experiments in fluids 41, 965–974 (2006).

11 H. Wagner, “Über stoß-und gleitvorgänge an der oberfläche von flüssigkeiten”, ZAMM-Journal of Applied Mathematics and Mechanics/Zeitschrift für Angewandte Mathematik und Mechanik 12, 193–215 (1932).

12 T. Von Karman, “The impact of seaplane floats during landing. NACA TN 321”, October, Washington (1929).

13 M. Jamali, A. Rostamijavanani, N. Nouri, and M. Navidbakhsh, “An experimental study of cavity and Worthington jet formations caused by a falling sphere into an oil film on water”, Applied Ocean Research 102, 102319 (2020).

14 J. T. Antolik, J. L. Belden, N. B. Speirs, and D. M. Harris, “Slamming forces during water entry of a simple harmonic oscillator”, Journal of Fluid Mechanics 974, A23 (2023).

15 F. C. Korkmaz and B. Güzel, “Water entry of cylinders and spheres under hydrophobic effects; Case for advancing deadrise angles”, Ocean engineering 129, 240–252 (2017).

16 M. Barjasteh, H. Zeraatgar, and M. J. Javaherian, “An experimental study on water entry of asymmetric wedges”, Applied Ocean Research 58, 292–304 (2016).

17 J. Qian, S. Zhang, and H. Jin, “Computer simulation of “splash control” and research of the rip entry technique in competitive diving”, International Journal of Sports Science and Engineering 4, 165–173 (2010).

18 L. Vincent, T. Xiao, D. Yohann, S. Jung, and E. Kanso, “Dynamics of water entry”, Journal of Fluid Mechanics 846, 508–535 (2018).

19 A. Pandey, J. Yuk, B. Chang, F. E. Fish, and S. Jung, “Slamming dynamics of diving and its implications for diving-related injuries”, Science advances 8, eabo5888 (2022).

20 B. Chang, M. Croson, L. Straker, S. Gart, C. Dove, J. Gerwin, and S. Jung, “How seabirds plunge-dive without injuries”, Proceedings of the National Academy of Sciences 113, 12006–12011 (2016).

21 E. Gregorio, E. Balaras, and M. C. Leftwich, “Air cavity deformation by single jointed diver model entry bodies”, Experiments in Fluids 64, 168 (2023).

22 J. G. Brown, L. D. Abraham, and J. J. Bertin, “Descriptive analysis of the rip entry in competitive diving”, Research Quarterly for Exercise and Sport 55, 93–102 (1984).

23 B. M. Currie, M. K. Drew, M. Hetherington, G. Waddington, N. A. Brown, and L. A. Toohey, “Diving into the health problems of competitive divers: a systematic review of injuries and illnesses in pre-elite and elite diving athletes”, Sports health, 19417381241255329 (2024).

24 E. Belilos, S. Jow, and M. Maxwell, “Descriptive epidemiology of high school swimming and diving injuries”, Clinical journal of sport medicine 33, 428–434 (2023).

25 J.-R. Delaloye, F. Sander, J. Murar, T. Tischer, and L. Ernstbrunner, “Water Jumping Sports”, Injury and Health Risk Management in Sports: A Guide to Decision Making, 651–657 (2020).

26 S. Gekle and J. M. Gordillo, “Generation and breakup of Worthington jets after cavity collapse. Part 1. Jet formation”, Journal of Fluid Mechanics 663, 293–330 (2010).

27 J. Glasheen and T. McMahon, “Vertical water entry of disks at low Froude numbers”, Physics of Fluids 8, 2078–2083 (1996).

28 J. M. Aristoff and J. W. Bush, “Water entry of small hydrophobic spheres”, Journal of Fluid Mechanics 619, 45–78 (2009).

29 S. Gekle, A. van der Bos, R. Bergmann, D. van der Meer, and D. Lohse, “Noncontinuous Froude number scaling for the closure depth of a cylindrical cavity”, Physical review letters 100, 084502 (2008).

30 A. May, “Vertical entry of missiles into water”, Journal of Applied Physics 23, 1362–1372 (1952).

31 G. Birkhoff and R. Isaacs, “Transient cavities in air–water entry”, Navord Rep 1490, 68–72 (1951).

32 D. Choi, J. Byun, and H. Park, “Analysis of liquid column atomization by annular dual-nozzle gas jet flow”, Journal of Fluid Mechanics 943, A25 (2022).

